# CYP1A1/1A2 enzymes mediate glucose homeostasis and insulin secretion in mice in a sex-specific manner

**DOI:** 10.1101/2024.02.16.580670

**Authors:** Ma. Enrica Angela Ching, Myriam P Hoyeck, Lahari Basu, Rayanna Merhi, Erin van Zyl, Angela M Crawley, Jennifer E Bruin

## Abstract

**Aims/hypothesis:** The aryl hydrocarbon receptor (AhR) pathway is involved in cellular responses to a broad range of external stressors, making it an excellent candidate for understanding the interaction between environmental factors and type 2 diabetes risk. Studies suggest deleting or downregulating AhR protects against metabolic dysfunction in high-fat diet (HFD) fed mice; however, the contribution of downstream AhR targets in driving this phenotype remains unexamined. Cytochrome P450 1A1 and 1A2 (CYP1A1/1A2) are canonical AhR targets that encode xenobiotic metabolism enzymes. Interestingly, we have demonstrated that HFD feeding increases *Cyp1a1* expression in mouse islets, which suggests CYP1A enzymes are involved in the response to metabolic stress. Since CYP1A1/1A2 activity can produce reactive oxygen intermediates, we hypothesized that chronic activation of these enzymes in tissues critical for regulating glucose homeostasis (e.g., liver, adipose, islets) will contribute to metabolic dysfunction following HFD feeding.

**Methods:** At 29 to 31 weeks of age, male and female global *Cyp1a1/1a2* knockout (*Cyp*^KO^) and wildtype littermate control (*Cyp*^WT^) mice were fed either a 45% HFD or standard rodent chow for 14 weeks. Metabolic assessments were conducted throughout the study.

**Results:** *Cyp*^KO^ females were partially protected from HFD-induced glucose intolerance compared to *Cyp*^WT^ females, but both genotypes exhibited similar levels of insulin resistance. *Cyp*^KO^ females also had lower plasma insulin levels *in vivo* and suppressed insulin secretion in isolated islets *ex vivo* compared to *Cyp*^WT^ females. Gene expression patterns in female islets were generally similar across genotype and diet groups. In contrast, *Cyp*^WT^ males became hyperinsulinemic and insulin resistant within 2 weeks of HFD feeding, while *Cyp*^KO^ males maintained normal plasma insulin levels and insulin sensitivity. HFD feeding upregulated *Cyp1a1* in *Cyp*^WT^ male islets and this was accompanied by elevation of other islet stress genes. Interestingly, HFD feeding did not induce these stress gene responses in *Cyp*^KO^ male islets, suggesting the islet stress response is mediated by activation of CYP1A1. We expected the global deletion of *Cyp1a1/1a2* to have pronounced effects in the liver, but surprisingly, changes in liver pathology were predominantly driven by diet and not genotype in both sexes. Similarly, overall adiposity and adipose tissue inflammation were not affected by genotype.

**Conclusions:** Our study highlights a novel role of islet *Cyp1a1/1a2* in shaping the systemic metabolic response to HFD feeding. Our data suggest that CYP1A1/1A2 enzymes are involved in glucose homeostasis, insulin secretion, and the islet stress response. Importantly, the effects of *Cyp1a1/1a2* deletion are sex-dependent.

## 1. Introduction

Over 500 million people are living with type 2 diabetes globally [1]. Recent estimates suggest the prevalence of type 2 diabetes could rise by another 50% and affect nearly 800 million people by 2045 [1]. Genetics, physical inactivity, and diet all contribute to type 2 diabetes risk, but increasing evidence suggests that other environmental factors, including pollutant exposure, also play a role [2–13]. The biological mechanisms underlying the relationship between environmental stressors and diabetes pathogenesis are complex and still poorly understood.

The aryl hydrocarbon receptor (AhR) recognizes a broad range of environmental stimuli and is best known for responding to xenobiotic exposures [14, 15]. However, AhR is also involved in other physiological processes, such as fat metabolism and inflammation [14, 15]. AhR is a ligand-activated transcription factor that interacts with both endogenous and exogenous compounds. Endogenous AhR ligands include tryptophan and heme metabolites [16, 17], while exogenous ligands include pollutants (e.g., dioxins, dioxin-like chemicals, polycyclic aromatic hydrocarbons) and food-derived compounds (e.g., polyphenols) [17, 18]. Upon ligand-binding, AhR translocates from the cytoplasm into the nucleus, where it forms a complex with the aryl hydrocarbon receptor nuclear translocator (ARNT) and binds to the promoter regions of its target genes [15].

Cytochrome P450 1a2 and 1a2 (*Cyp1a1/1a2*) are canonical AhR target genes that encode phase I xenobiotic metabolism enzymes [15]. These membrane-associated proteins are typically located in the mitochondria and endoplasmic reticulum of mammalian cells [18, 19]. CYP1A1/1A2 substrates include environmental pollutants, pharmaceutical drugs, and endogenous compounds (e.g., melatonin and estradiol) [18, 20]. CYP1A1/1A2 enzymes facilitate xenobiotic clearance by increasing the polarity of substrates through oxidation, allowing for further enzyme degradation [18]. However, CYP1A1/1A2 enzymes can also generate reactive oxygen species (ROS) and activate procarcinogens [20–23]. Despite their essential role in detoxification, CYP1A1/1A2 activity can have harmful effects.

Numerous studies have shown that downregulating or deleting AhR protects against high-fat diet (HFD)-induced obesity and glucose dysregulation in mice [24–31]. These studies emphasize the importance of the AhR pathway in metabolic adaptation to HFD, but the role of downstream AhR target genes, *Cyp1a1* and *Cyp1a2,* remains unclear. Furthermore, the focus of previous studies is typically on adipose- and liver-related outcomes, but our lab has previously shown that HFD feeding can upregulate *Cyp1a1* expression in the mouse endocrine pancreas (i.e., islets) [32]. Although islets are critical for maintaining glucose homeostasis, the contribution of AhR signalling to islet function and adaptation in response to HFD has not been well characterized.

To address these gaps, we performed a 14-week HFD feeding study with global *Cyp1a1/1a2* double knockout mice (*Cyp*^KO^) and wildtype littermate controls (*Cyp*^WT^) of both sexes. We tracked body weight, adiposity, and fasting blood glucose throughout the study, and assessed systemic glucose tolerance, insulin resistance, and plasma insulin levels. We also evaluated liver and adipose phenotypes alongside islet-specific outcomes, such as measurements of glucose-stimulated insulin secretion (GSIS) *ex vivo* and islet gene expression, at the study endpoint.

## 2. Materials and Methods

### 2.1 Animals

Male and female global *Cyp*^KO^ and *Cyp*^WT^ littermate controls (C57BL/6J background) were bred and housed in the Carleton University vivarium. *Cyp*^KO^ mice were initially generated by Dr. Daniel Nebert’s group [33] and were kindly provided to us by Dr. Frank Gonzalez (University of Cincinnati). All mice were kept on a 12-hour light/dark cycle and had *ad libitum* access to standard rodent chow (Harlan Laboratories, Teklad Diet #2018, Madison, WI, USA).

Experimental mice were 29 to 31 weeks old at the start of the study **(Figure 1A)**. The mice in each diet group were matched based on body weight and fasting blood glucose values obtained from week -5 to week 0. At week 0, a subset of mice was transferred to a 45% HFD (Research Diets D12451, New Brunswick, NJ, USA), or remained on standard rodent chow for 14 weeks. There were 7-8 mice per group for females (n_females_ = 8 *Cyp*^WT^-Chow, 8 *Cyp*^WT^-HFD, 7 *Cyp*^KO^-Chow, 8 *Cyp*^KO^-HFD) and 5-8 mice per group for males (n_males_ = 8 *Cyp*^WT^-Chow, 6 *Cyp*^WT^-HFD, 5 *Cyp*^KO^-Chow, 5 *Cyp*^KO^-HFD).

**Figure 1.**
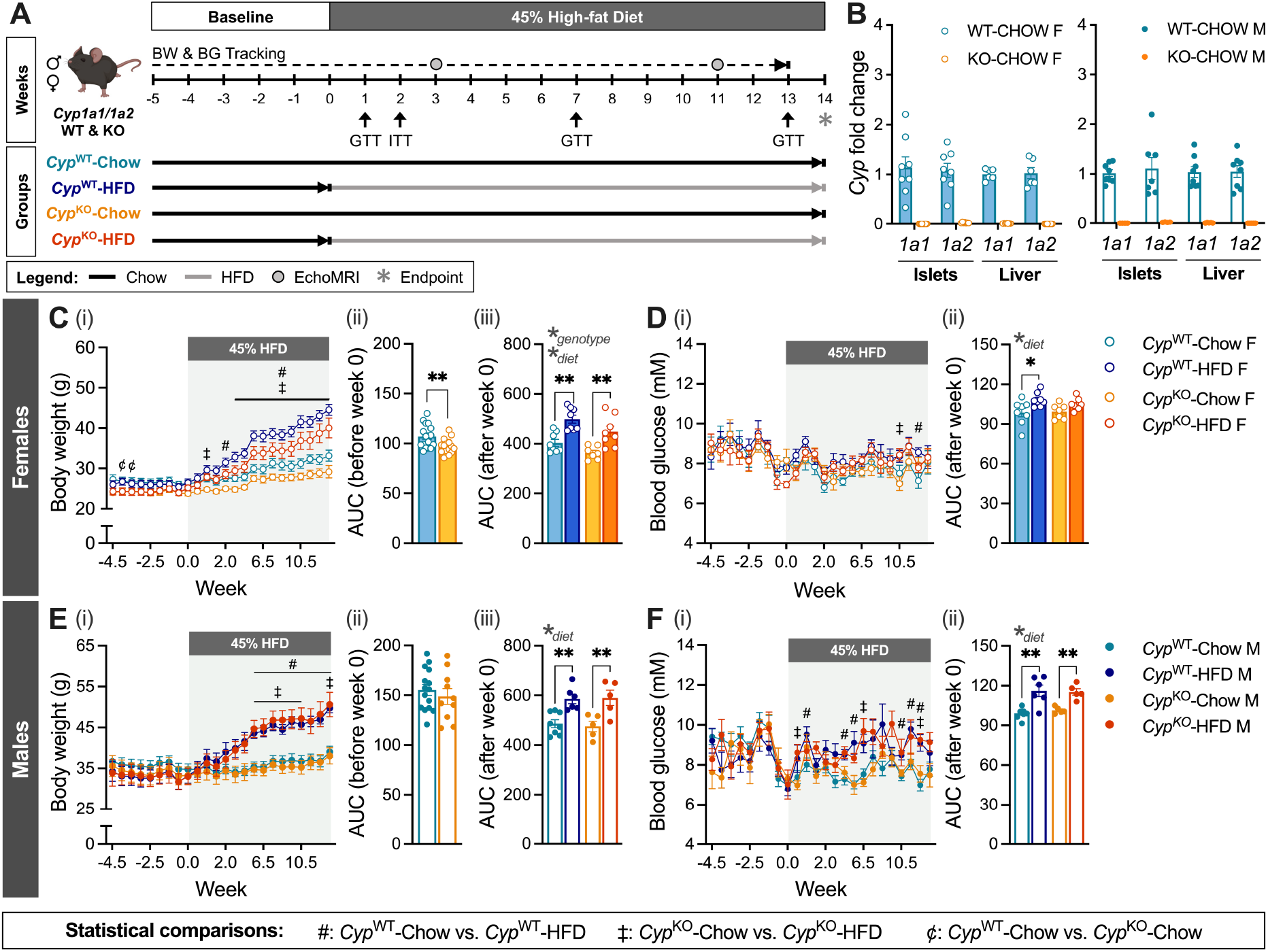
*Cyp*^KO^ females had lower body weight than *Cyp*^WT^ females on a chow diet. (A) Male and female *Cyp1a1/1a2* knockout (*Cyp*^KO^) and wildtype (*Cyp*^WT^) mice were tracked for 20 weeks, including 6 weeks of baseline tracking (weeks -5 to 0). At week 0, half of the mice were transferred to 45% high-fat diet (HFD) or remained on chow diet for another 14 weeks (n = 7-8 mice per group for females, n = 5-8 mice per group for males). Mice were euthanized at week 14 for tissue collection. BW = body weight, BG = blood glucose, GTT = glucose tolerance test. (B) *Cyp1a1* and *Cyp1a2* gene expression in islets and liver at week 14 in chow-fed male and female mice. (C, E) Body weight in (C) female and (E) male mice, presented as (i) a line graph, (ii) area under the curve (AUC) of BW prior to week 0, and (iii) AUC of BW data from week 0 to 13. (D, F) Fasting blood glucose in (D) female and (F) male mice, presented as (i) a line graph, and (ii) AUC of BG data from weeks 0 to 13. All data are presented as mean ± SEM. Individual data points on bar graphs represent biological replicates (different mice). The following statistical tests were used: (C-F, i) line graphs: mixed-effects analysis with Tukey’s post-hoc test at p < 0.05, comparison groups – #: *Cyp*^WT^-Chow vs. *Cyp*^WT^-HFD, ‡: *Cyp*^KO^-Chow vs. *Cyp*^KO^-HFD, ¢: *Cyp*^WT^-Chow vs. *Cyp*^KO^-Chow; (Cii-iii, Eii-iii, Dii, Fii); bar graphs: 2-way ANOVA with Tukey’s post-hoc test, *p < 0.05, **p < 0.01.

At week 14, all mice were euthanized to isolate pancreatic islets and collect liver and peripancreatic fat tissues. Islets were used for *ex vivo* GSIS assays and quantitative real-time PCR (qPCR); liver and peripancreatic fat tissues were used for histology and qPCR. Islets for qPCR were stored in buffer RLT (RNeasy Micro Kit, QIAGEN, Cat # 74004) + 1% β-Mercaptoethanol (Sigma Aldrich, Cat # M3148) and kept at -80°C. Liver and peripancreatic fat tissues for histology were stored in 4% paraformaldehyde (PFA; Fisher Scientific, Cat # AAJ19943K2) at 4°C overnight, then transferred to 70% ethanol for long-term storage at 4°C until paraffin embedding; tissues for qPCR were flash-frozen immediately after collection and stored at -80°C. All experiments were performed in accordance with guidelines set by Carleton University’s Animal Care Committee and the Canadian Council on Animal Care.

### 2.2 *In vivo* metabolic assessments

Body weight and fasting blood glucose levels were measured after a 4- to 6-hour morning fast, once or twice per week throughout the study. Blood glucose measurements were taken using handheld glucometers, specifically the OneTouch Verio Flex (LifeScan Canada, measuring range: 1.1 mmol/L to 33.3 mmol/L) for weeks -5 to -1 of the study, and the MediSure Multi-Patient Use Glucose Meter (MediSure Canada, measuring range: 1.1 mmol/L to 41.7 mmol/L) for weeks 0 to 13 of the study. Whole body fat composition was assessed using an EchoMRI-700 (EchoMRI LLC, Houston, TX, USA) at weeks 3 and 11 of the study **(Figure 1A)**.

Glucose tolerance tests (GTTs) were performed following a 6-hour morning fast. Blood glucose levels were measured at t = 0, 15, 30, 60, and 90 minutes post-glucose injection using a handheld glucometer. Saphenous blood was collected at t = 0, 15, 30, and 60 minutes during GTTs to measure plasma insulin levels by ELISA (ALPCO rodent ultrasensitive insulin ELISA, Cat # 80-INSMSU-E10, Salem, NH, USA). Mice received 1.5 g/kg of glucose (DMVet Dextrose 50%, DIN: 02420880) for all GTTs except the GTT at week 1, where they received 2 g/kg of glucose. The glucose dose was reduced to avoid exceeding the detection limit of the glucometer. We performed an intraperitoneal GTT (ipGTT) at weeks 1 **(Figure 2A, 2D, 3A, 4A)** and 7 **(Figure 2B, 2E, 3B, 4B)**, and an oral GTT (oGTT) at week 13 **(Figure 2C, 2F, 3C, 4C)**, wherein the mice received glucose by oral gavage.

**Figure 2.**
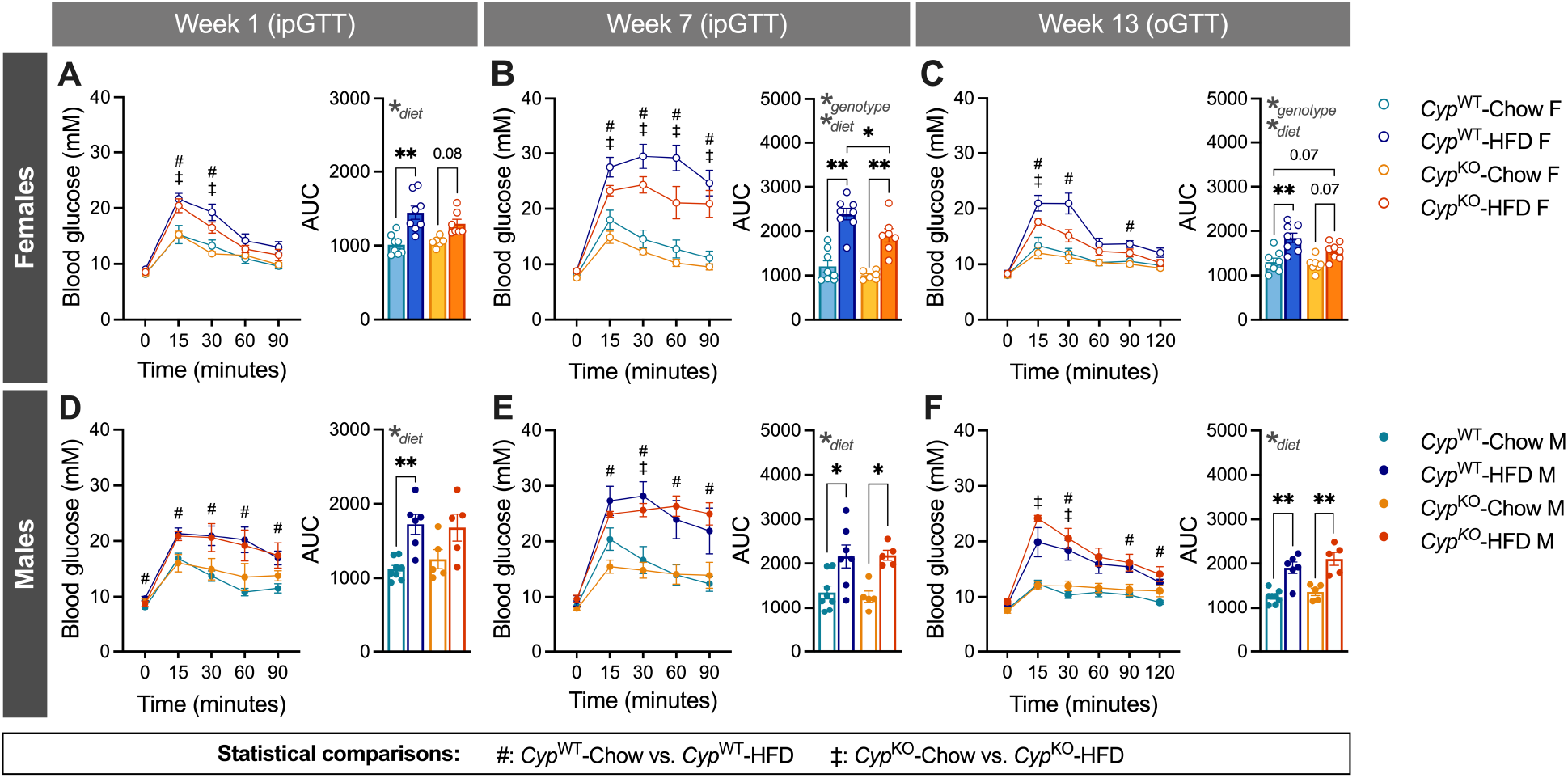
*Cyp*^KO^ females developed less severe glucose intolerance on HFD compared to *Cyp*^WT^ females. Glucose tolerance (GTT) was assessed at (A, D) week 1 and (B, E) week 7 with an intraperitoneal GTT (ipGTT), and at (C, F) week 13 with an oral GTT (oGTT) in (A-C) female and (D-F) male *Cyp*^WT^ and *Cyp*^KO^ mice (see Figure 1A for the study timeline). Mice received a glucose dose of 2 g/kg at (A, D) week 1, and 1.5 g/kg at (B, E) week 7 and (C, F) week 13. Data is presented as a line graph and area under the curve (AUC). All data are presented as mean ± SEM. Individual data points on AUC graphs represent biological replicates (different mice). The following statistical tests were used: line graphs: mixed-effects analysis with Tukey’s post-hoc test at p < 0.05, comparison groups – #: *Cyp*^WT^-Chow vs. *Cyp*^WT^-HFD, ‡: *Cyp*^KO^-Chow vs. *Cyp*^KO^-HFD; bar graphs: 2-way ANOVA with Tukey’s post-hoc test, *p < 0.05, **p < 0.01.

**Figure 3.**
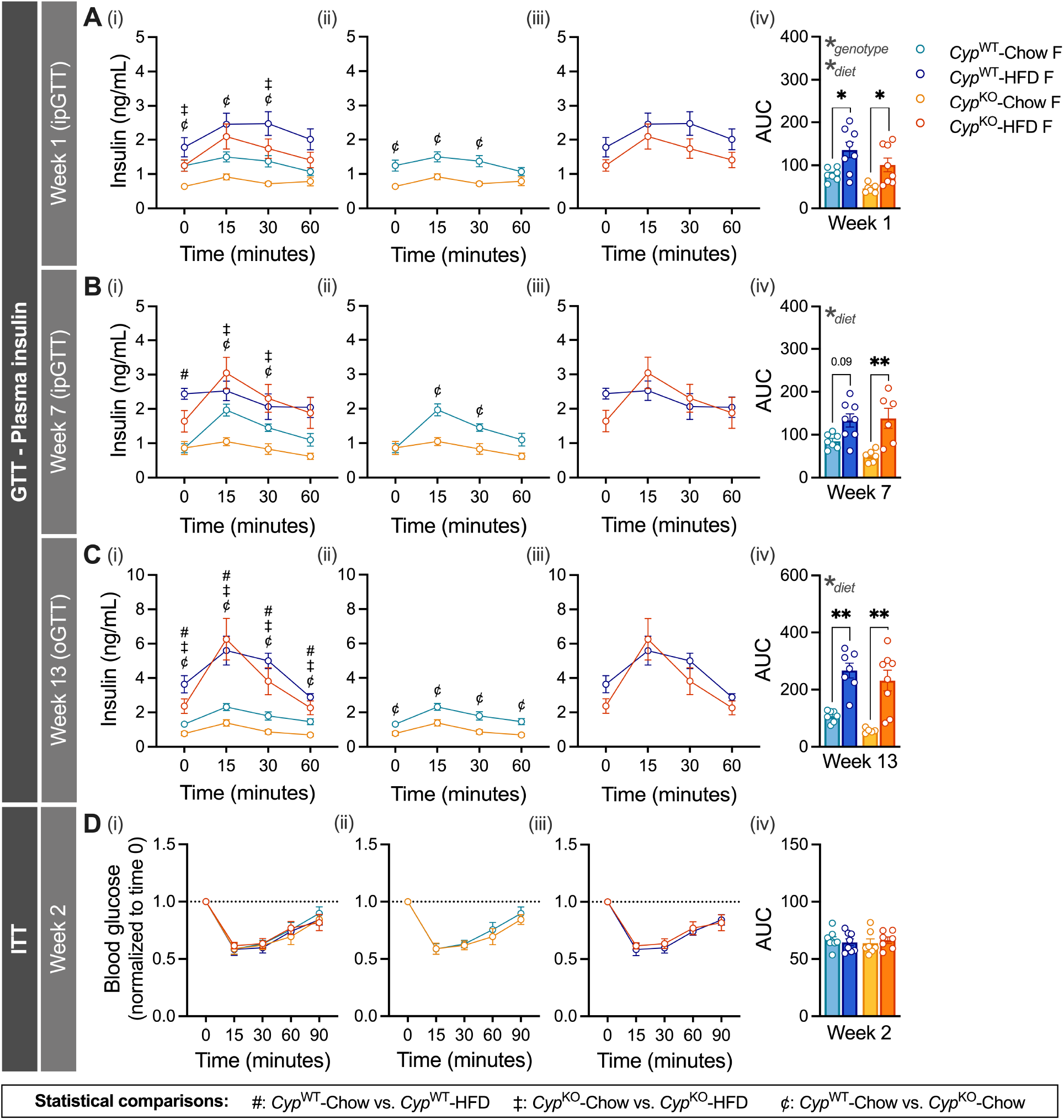
*Cyp*^KO^ female mice had lower plasma insulin levels than *Cyp*^WT^ females on a chow diet. Glucose tolerance tests (GTT) were performed at weeks 1, 7, and 13, and an insulin tolerance test (ITT) was performed at week 2 (see Figure 1A for the study timeline). Plasma insulin levels during the GTT at (A) week 1, (B) week 7, and (C) week 13 in female *Cyp*^WT^ and *Cyp*^KO^ mice. Mice received either (A) 2 g/kg glucose or (B, C) 1.5 g/kg of glucose (n = 5-8 per group). Glucose was administered (A, B) intraperitoneally (ipGTT), or (C) orally (oGTT). (D) Blood glucose levels during the ITT; mice received a dose of 1.0 IU/kg of insulin by intraperitoneal injection (n = 7-9 per group). Blood glucose values for the ITT were normalized to baseline levels at t = 0. Data is presented as (i-iii) line graphs showing (i) all groups, (ii) chow-fed mice only, (iii) HFD-fed mice only, and (iv) area under the curve (AUC). All data are presented as mean ± SEM. Individual data points on AUC graphs represent biological replicates (different mice). The following statistical tests were used: (A-D, i-iii) line graphs: mixed-effects analysis with Tukey’s post-hoc test at p < 0.05, comparison groups – #: *Cyp*^WT^-Chow vs. *Cyp*^WT^-HFD, ‡: *Cyp*^KO^-Chow vs. *Cyp*^KO^-HFD, ¢: *Cyp*^WT^-Chow vs. *Cyp*^KO^-Chow. (A-D, iv). Bar graphs: 2-way ANOVA with Tukey’s post-hoc test; *p < 0.05, **p < 0.01.

**Figure 4.**
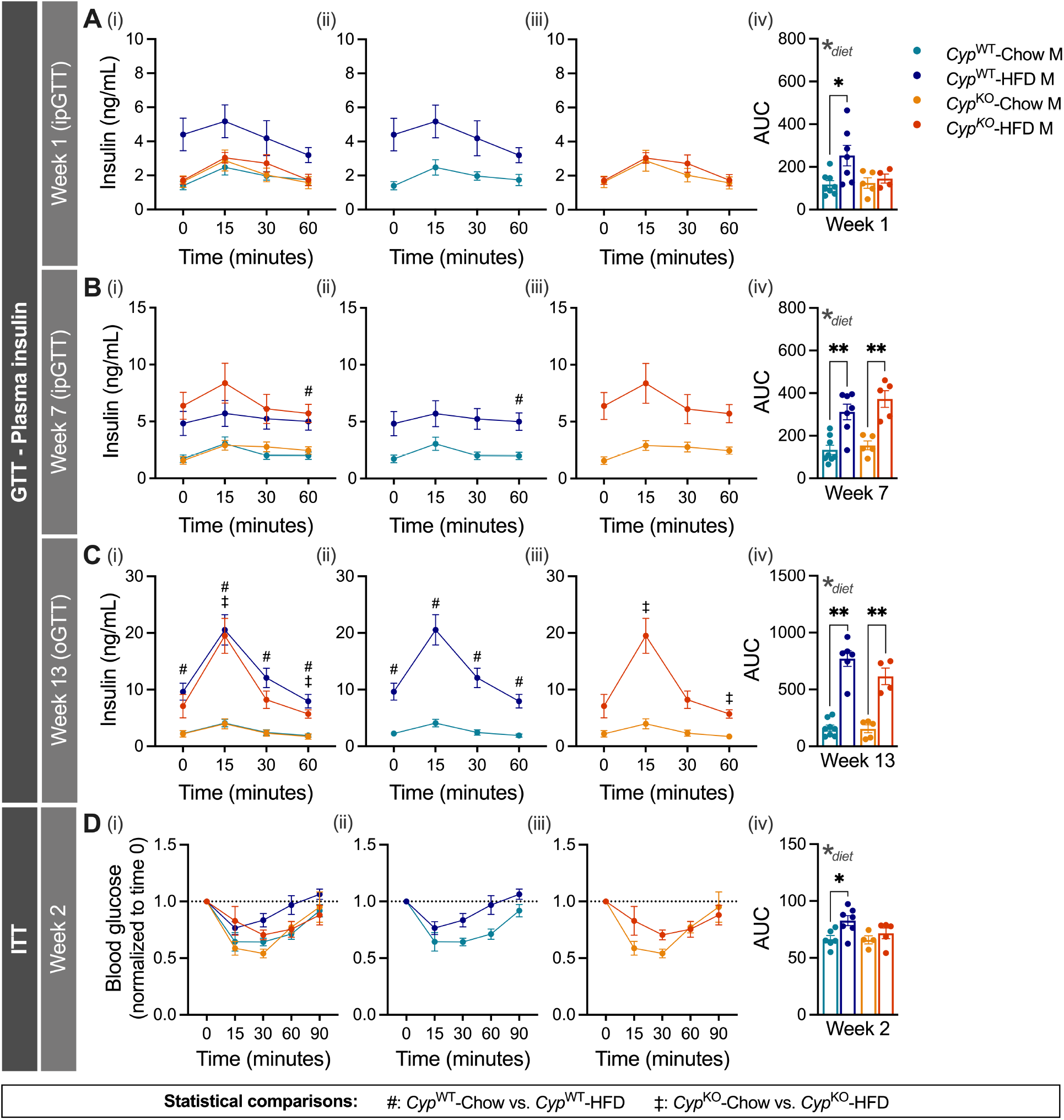
HFD-fed *Cyp*^KO^ males became hyperinsulinemic later than HFD-fed *Cyp*^WT^ males. Glucose tolerance tests (GTT) were performed at weeks 1, 7, and 13, and an insulin tolerance test (ITT) was performed at week 2 (see Figure 1A for the study timeline). Plasma insulin levels during the GTT at (A) week 1, (B) week 7, and (C) week 13 in male *Cyp*^WT^ and *Cyp*^KO^ mice. Mice received either (A) 2 g/kg glucose or (B, C) 1.5 g/kg of glucose (n = 4-8 per group). Glucose was administered (A, B) intraperitoneally (ipGTT), or (C) orally (oGTT). (D) Blood glucose levels during the ITT; mice received a dose of 1.0 IU/kg of insulin by intraperitoneal injection (n = 4-7 per group). Blood glucose values for the ITT were normalized to baseline levels at t = 0. Data is presented as (i-iii) line graphs showing (i) all groups, (ii) WT mice only, (iii) KO mice only, and (iv) area under the curve (AUC). All data are presented as mean ± SEM. Individual data points on AUC graphs represent biological replicates (different mice). The following statistical tests were used: (A-D, i-iii) line graphs: mixed-effects analysis with Tukey’s post-hoc test at p < 0.05, comparison groups – #: *Cyp*^WT^-Chow vs. *Cyp*^WT^-HFD, ‡: *Cyp*^KO^-Chow vs. *Cyp*^KO^-HFD, ¢: *Cyp*^WT^-Chow vs. *Cyp*^KO^-Chow. (A-D, iv). Bar graphs: 2-way ANOVA with Tukey’s post-hoc test; *p < 0.05, **p < 0.01.

For the insulin tolerance test (ITT), mice received an intraperitoneal injection of 1.0 IU/kg insulin (Novolin^®^ge Toronto, Novo Nordisk Canada, DIN: 02024233) after a 4-hour morning fast. We measured blood glucose levels at t = 0, 15, 30, 60, and 90 minutes.

2.3 Mouse islet isolation

We performed pancreatic duct injections with collagenase (Sigma-Aldrich, Cat # C7657) dissolved in Hanks’ balanced salt solution (HBSS: 137 mM NaCl, 5.4 mM KCl, 4.2 mM NaH_2_PO4, 4.1 mM KH_2_PO_4_, 10 mM HEPES, 1 mM MgCl_2_, 5 mM dextrose, pH 7.2). The inflated pancreas tissues were transferred to cold HBSS and kept on ice until further processing. To resume digestion, the pancreas samples were incubated at 37°C for 10-11 minutes, then vigorously shaken. Cold HBSS with 1 mM CaCl_2_ was immediately added afterwards to stop digestion. The samples were then washed three times in cold HBSS with CaCl_2_. After the third wash, the islets were resuspended in RPMI media (Wisent, Cat # 350-000-CL) supplemented with 1% Penicillin-Streptomycin (Gibco, Cat # 15-140-122) and 10% FBS (Sigma-Aldrich, Cat # F1051). The suspension was then filtered and hand-picked to >95% purity using dissecting microscopes (Zeiss Stemi 508). Islets were incubated overnight at 37°C and 5% CO_2_ in supplemented RPMI to allow for recovery prior to conducting experiments. Isolated islets were used for an *ex vivo* GSIS assay or stored in buffer RLT + 1% β-Mercaptoethanol and kept at -80°C for qPCR.

### 2.4 *Ex vivo* GSIS assay

To assess beta cell function, we performed an *ex vivo* GSIS assay. We used 25 islets per replicate (n = 3 technical replicates per mouse, n = 4-8 mice per experimental group). Islets were hand-picked into microcentrifuge tubes and washed with warm Krebs-Ringer Bicarbonate HEPES (KRBB) buffer (0 mM glucose, 115 mM NaCl, 5 mM KCl, 2.5 mM CaCl_2_, 1 mM MgCl_2_, 10 mM HEPES, 24 mM NaHCO_3_, 0.1 % w/v BSA). Islets were pre-incubated in 500 µL of low glucose KRBB solution (LG; 2.8 mM; Sigma-Aldrich, Cat # G8769-100mL) for 1 hour at 37°C and 5% CO_2_ and the supernatant was discarded. Islets were then incubated for 1 hour in 500 µL of LG, followed by 1 hour in 500 µL of high glucose KRBB solution (HG; 16.7 mM), and lastly in acid ethanol (AE; 1.5% vol./vol. HCl in 70% vol./vol. ethanol) overnight at 4°C to measure total insulin content. Supernatants were collected following each incubation. For AE samples, 450 µL of the supernatant was collected and neutralized with 450 µL of 1M Tris base. All GSIS samples were kept at - 30°C for long-term storage. Insulin concentrations were measured using the rodent insulin chemiluminescent ELISA kit (ALPCO, Cat # 80-INSMR-CH10) according to the manufacturer’s instructions. Stimulation index was calculated for each biological replicate as follows: [average insulin concentration in HG-KRBB] / [average insulin concentration in LG/KRBB].

### 2.5 Histology

PFA-fixed liver and peripancreatic fat samples were embedded, processed, and stained with hematoxylin and eosin stain (H&E) and Masson’s trichome stain (MT) at the Louise Pelletier Histology Core Facility at the University of Ottawa. A pathologist with Nour Histopathology Consultation Services (Ottawa, ON, Canada) evaluated MT-stained liver samples for fibrosis, and H&E-stained liver samples for steatosis, ballooning, and inflammation. Fibrosis scoring was based on the METAVIR scoring system, which ranges from 0 to 4, with 0 representing no fibrosis and 4 representing cirrhosis [34]. Scoring for steatosis, ballooning, and inflammation were based on the non-alcoholic steatohepatitis (NASH) grading system developed by the NASH Clinical Research Network [35]. Each category (i.e., steatosis, ballooning, inflammation) was scored separately, with scores ranging from 0 to 3 (0 = no observable disease phenotype, 1 = mild, 2 = moderate, 3 = severe) [35].

Peripancreatic fat samples were embedded at the Louise Pelletier Histology Core Facility and tissue sections were stained with H&E (Epredia™ Modified Harris Hematoxylin, Fisher Scientific, Cat # 22-050-206; Eosin Y disodium salt, Sigma-Aldrich, Cat # E4382) in the Bruin Lab. Slides were imaged using a ZEISS Axio Observer 7 microscope with ZEN 2.6 (blue edition) software. Adipose inflammation in H&E-stained tissues was graded from 0 to 3 based on previously described qualitative criteria [36]. A score of 0 means there was no visible inflammation or inflammatory cells were scattered and rarely observed; a score of 1 represents minimal local inflammation; a score of 2 represents mild diffuse inflammation; and a score of 3 represents moderate to severe diffuse inflammation. All adipose slides were evaluated and scored by a single evaluator who was blinded to the treatment conditions.

### 2.6 Quantitative real-time PCR

RNA was extracted from mouse islets using the RNAeasy Micro Kit (QIAGEN, Cat # 74004); RNA was extracted from flash-frozen liver and peripancreatic fat tissues using TRIzol^TM^ (Invitrogen, #15596018; Carlsbad, CA, USA). cDNA was made using the iScript gDNA Clear cDNA Synthesis Kit with DNase treatment (Bio-Rad, Cat # 1725035). RNA extraction and cDNA synthesis were conducted following the manufacturers’ instructions.

Relative mRNA levels in islets, liver, and peripancreatic fat were measured by qPCR using the SsoAdvanced Universal SYBR Green Supermix (Bio-Rad, Cat # 1725271) and Precision Blue Real-Time PCR Dye (Bio-Rad, Cat # 1725275). The qPCR reactions were run on a CFX384 qPCR detection system (Bio-Rad, Cat # 43174). Primer sequences are in **Supplement Table 1**. Primers were ordered from Integrated DNA Technologies (IDT, USA). The 2^-ΔΔCT^ method was used to analyze the qPCR data, and *PPIA* or rRNA was used as the reference gene since these genes showed stable expression across all groups.

### 2.7 Statistical Analyses

Statistical analyses were performed using GraphPad Prism (Version 10.0.1 for macOS). For analyses assessing the effects of diet and genotype over time (e.g., GTT, ITT), a repeated measures two-way ANOVA or a mixed-effects model with Geisser-Greenhouse correction was used followed by Tukey’s post-hoc for multiple comparisons. For all other analyses (e.g., area under the curve (AUC), qPCR), a two-way ANOVA with Tukey’s post-hoc was used. Statistical significance was considered at p < 0.05. Data are presented as mean ± SEM.

## 3. Results

### 3.1 *Cyp*^KO^ females had lower body weight than *Cyp*^WT^ females on a chow diet

We first validated our global knockout model by measuring *Cyp1a1* and *Cyp1a2* levels in isolated islets and liver tissues. Compared to *Cyp*^WT^ mice, levels of *Cyp1a1* or *Cyp1a2* were not detectable in *Cyp*^KO^ islets and liver tissues **(Figure 1B)**. We next measured fasting blood glucose and body weight of *Cyp*^WT^ and *Cyp*^KO^ mice on chow diet for 6 weeks (weeks -5 to 0) followed by HFD for 14 weeks. During the baseline tracking period, chow-fed *Cyp*^KO^ female mice had significantly lower body weight than *Cyp*^WT^ females (**Figure 1Ci, ii**). After switching to HFD, genotype still accounted for some of the variations in body weight (12%), but this was outsized by the effect of diet, which explained 47% of the total variation. Within each diet group (chow vs HFD), *Cyp*^KO^ females were no longer significantly smaller than *Cyp*^WT^ females, which may be due to the reduction in sample size and statistical power **(Figure 1Ciii)**. Fasting blood glucose levels were similar between *Cyp*^WT^ and *Cyp*^KO^ females throughout the study, irrespective of diet **(Figure 1D)**. There were no genotype-based differences in body weight **(Figure 1E)** or fasting blood glucose **(Figure 1F)** in males fed a chow or HFD diet.

### 3.2 *Cyp*^KO^ females developed less severe glucose intolerance on HFD compared to *Cyp*^WT^ females

There were no significant differences in glucose tolerance between *Cyp*^WT^ and *Cyp*^KO^ females fed a chow diet (**Figure 2A-C**). However, HFD-fed *Cyp*^KO^ females developed less severe glucose intolerance compared to *Cyp*^WT^ females at weeks 7 and 13 **(Figure 2B-C)**. The results from the ipGTT at week 7 **(Figure 2B)** were replicated in the oGTT at week 13 **(Figure 2C)**, which suggests that the improved glucose tolerance of HFD-fed *Cyp*^KO^ females is not dependent on incretin hormones. In contrast to females, differences in glucose tolerance in males were driven entirely by diet. *Cyp*^WT^ and *Cyp*^KO^ males had similar glucose tolerance on either chow or HFD **(Figure 2D-F)**. Altogether, our data indicate that the presence of *Cyp1a1/1a2* is partially driving glucose intolerance in HFD-fed female mice.

### 3.3 *Cyp*^KO^ female mice had lower plasma insulin levels than *Cyp*^WT^ females on a chow diet

To further assess glucose homeostasis, we measured plasma insulin levels during the GTTs and insulin sensitivity during an ITT. Surprisingly, chow-fed *Cyp*^KO^ females consistently had lower plasma insulin levels during GTTs than chow-fed *Cyp*^WT^ females **(Figure 3A-C, ii)**, but there were no differences in plasma insulin levels between HFD-fed *Cyp*^WT^ and *Cyp*^KO^ females **(Figure 3A-C, iii)**. Although *Cyp*^KO^-Chow females required lower plasma insulin levels than *Cyp*^WT^-Chow females to maintain similar blood glucose levels during a GTT **(Figure 2A-C)**, *Cyp*^KO^-Chow females were not more insulin sensitive than *Cyp*^WT^-Chow females **(Figure 3D)**. Insulin tolerance was also unchanged in *Cyp*^KO^-HFD compared to *Cyp*^WT^-HFD females **(Figure 3D)**.

### 3.4 HFD-fed *Cyp*^KO^ males became hyperinsulinemic later than HFD-fed *Cyp*^WT^ males

*Cyp*^KO^ males displayed a delayed onset of HFD-induced hyperinsulinemia **(Figure 4A-C, i, iii, iv)** compared to *Cyp*^WT^ males. After 1 week of HFD feeding, *Cyp*^WT^-HFD males developed pronounced hyperinsulinemia **(Figure 4Ai, ii, iv)**, whereas *Cyp*^KO^-HFD males exhibited normal plasma insulin levels compared to chow-fed males **(Figure 4Ai, iii, iv)**. By weeks 7 and 13, both HFD-fed *Cyp*^WT^ and *Cyp*^KO^ males were hyperinsulinemic compared to their chow-fed counterparts **(Figure 4B, C)**. At week 2, *Cyp*^WT^-HFD males were also modestly insulin resistant compared to *Cyp*^WT^-Chow males, whereas *Cyp*^KO^-HFD had similar insulin sensitivity to *Cyp*^KO^-Chow males **(Figure 4D)**. These data suggest *Cyp1a1/1a2* are involved in promoting HFD-induced hyperinsulinemia and insulin resistance in male mice.

### 3.5 Liver and adipose tissues were affected by HFD feeding but not genotype

We assessed liver pathology and gene expression at the study endpoint (week 14) to examine if peripheral changes in the liver could explain the metabolic differences between HFD-fed *Cyp*^WT^ and *Cyp*^KO^ mice. Differences in liver pathology were driven by HFD feeding and not by genotype in both male and female mice **(Figure 5A, C, E-F)**; *Cyp*^WT^ and *Cyp*^KO^ mice developed similar liver fibrosis severities, steatosis, ballooning, and inflammation **(Figure 5A, C)**. As expected, *Cyp1a1* and *Cyp1a2* expression were absent in the liver of *Cyp*^KO^ mice **(Figure 5Bi-ii, Di-ii)**. While liver *Cyp1a1* levels were unaffected by diet, *Cyp1a2* expression was significantly lower in HFD-fed *Cyp*^WT^ mice compared to chow-fed *Cyp*^WT^ mice, irrespective of sex **(Figure 5Bii, Dii)**. In females, there was an overall effect of diet to decrease liver inflammation scores **(Figure 5C),** and this was also reflected in decreased levels of an inflammatory marker, *Tnfα*, in the liver of HFD-fed females **(Figure 5Biii)**. In males, there was an overall effect of diet to increase liver *Tnfα* levels **(Figure 5Diii)**, but no effect of diet on liver inflammation as determined by histopathology **(Figure 5C)**. Lastly, neither diet nor genotype affected the expression of *Slc2a2*, a major glucose transporter in liver **(Figure 5Bii-iii, Dii-iii)**.

**Figure 5.**
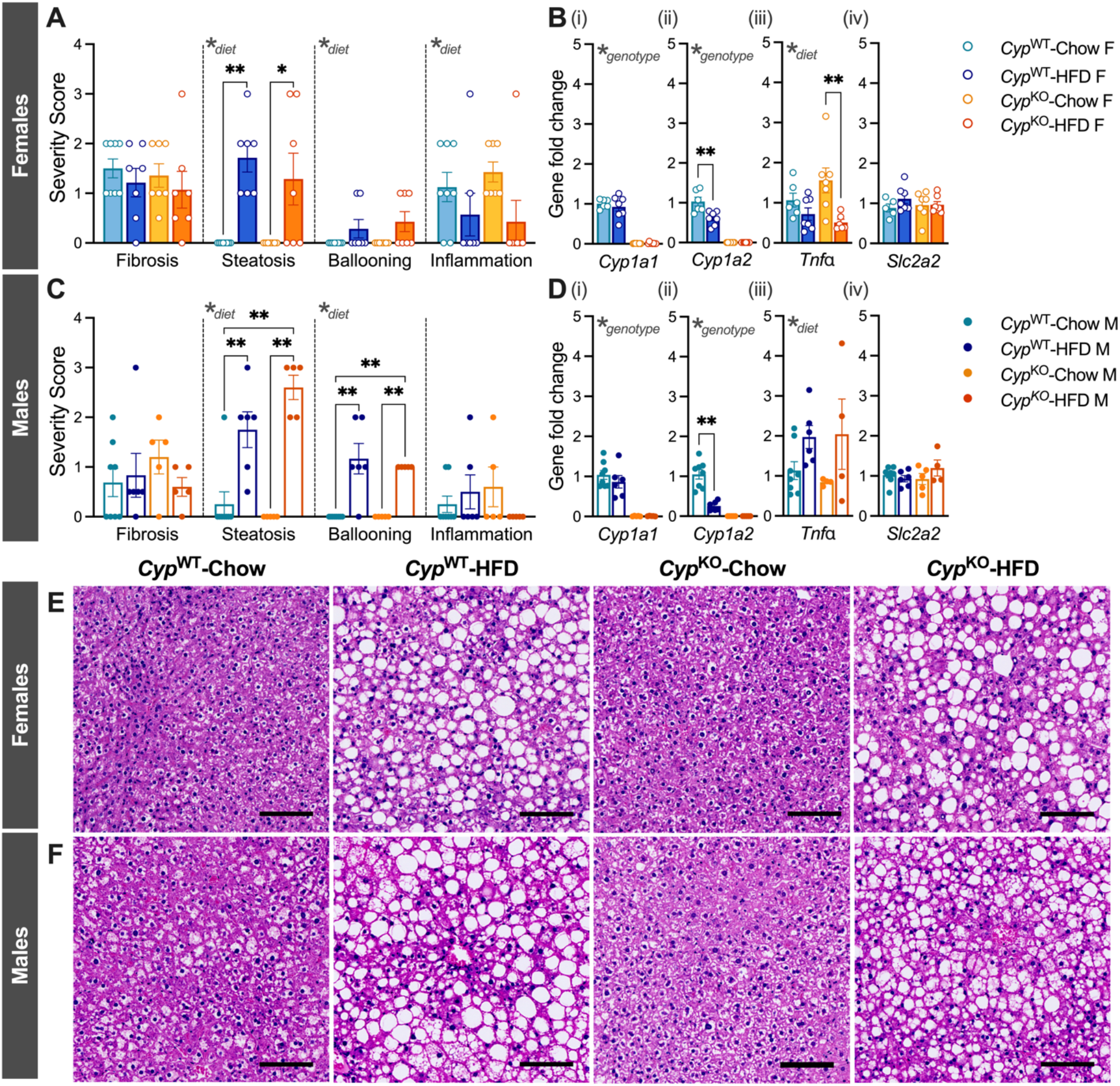
High-fat diet feeding induced liver damage in both *Cyp*^WT^ and *Cyp*^KO^ mice and reduced liver *Cyp1a2* expression in *Cyp*^WT^ mice. Liver samples were collected at week 14 for histology and qPCR analysis (see Figure 1A for the study timeline). (A, C) Liver damage severity scores and (E, F) representative H&E-stained liver images (20x magnification, scale bar = 100 µM) in (A, E) female (n = 7-8 mice per group) and (C, F) male mice (n = 5-8 mice per group). (A) Fibrosis scores range from 0 (no fibrosis) to 4 (cirrhosis); scores for steatosis, ballooning, and inflammation, range from 0 to 3 (0 = no observable disease phenotype, 1 = mild, 2 = moderate, 3 = severe). (B, D) *Cyp1a1*, *Cyp1a2, Tnf*α, and *Slc2a2* gene expression in liver from (B) female (n = 5-8 mice per group) and (D) male mice (4-8 mice per group). H&E: Hematoxylin and eosin stain. All data are presented as mean ± SEM. Individual data points on bar graphs represent biological replicates (different mice). The following statistical tests were used: 2-way ANOVA with Tukey’s post-hoc test; *p < 0.05, **p < 0.01.

To assess the effect of deleting *Cyp1a1/1a2* in adipose tissue, we measured % fat mass by EchoMRI at weeks 3 and 11 **(Supplement Figure 1A, D)**, adipose inflammation by histology at week 14 **(Supplement Figure 1B, E)**, and gene expression in peripancreatic fat by qPCR at week 14 **(Supplement Figure 1C, F)**. Changes in % fat mass were driven by diet in both sexes **(Supplement Figure 1A, D)**, and the severity of adipose inflammation was not affected by diet or genotype in either sex **(Supplement Figure 1B, E)**. There were also no significant differences in adipose *Tnfα* levels **(Supplement Figure 1Ci, Fi)** or *Slc2a4,* a glucose transporter gene prominent in adipose tissue, across genotype and diet groups **(Supplement Figure 1Cii, Fii)**.

In summary, the genotype-based differences in (1) glucose tolerance **(Figure 2A-C)** and plasma insulin levels in females **(Figure 3A-C)** and (2) the onset of HFD-induced hyperinsulinemia **(Figure 4A-C)** and insulin resistance **(Figure 4D)** in males were likely not explained by changes in liver pathology or adiposity. We next characterized the phenotype of pancreatic islets, which are critical for regulating glucose homeostasis. We assessed islet function and gene expression, but unfortunately did not have enough pancreas tissue for histology.

### 3.6 *Cyp1a1/1a2* are involved in regulating GSIS in both sexes

To assess islet function, we measured GSIS *ex vivo* in islets isolated at the study endpoint. Islets from *Cyp*^KO^ females showed a slightly reduced stimulation index compared to islets from *Cyp*^WT^ females, irrespective of diet **(Figure 6B)**. In males, islets from HFD-fed *Cyp*^WT^ mice had significantly lower stimulation index than islets from chow-fed *Cyp*^WT^ mice, but HFD feeding did not significantly impair the stimulation index of *Cyp*^KO^ male islets **(Figure 6E)**. There were no genotype- or diet-based differences in the concentration of secreted insulin following LG or HG incubations **(Figure 6A, D)** or total insulin content **(Figure 6C, F)** in islets from either sex.

**Figure 6.**
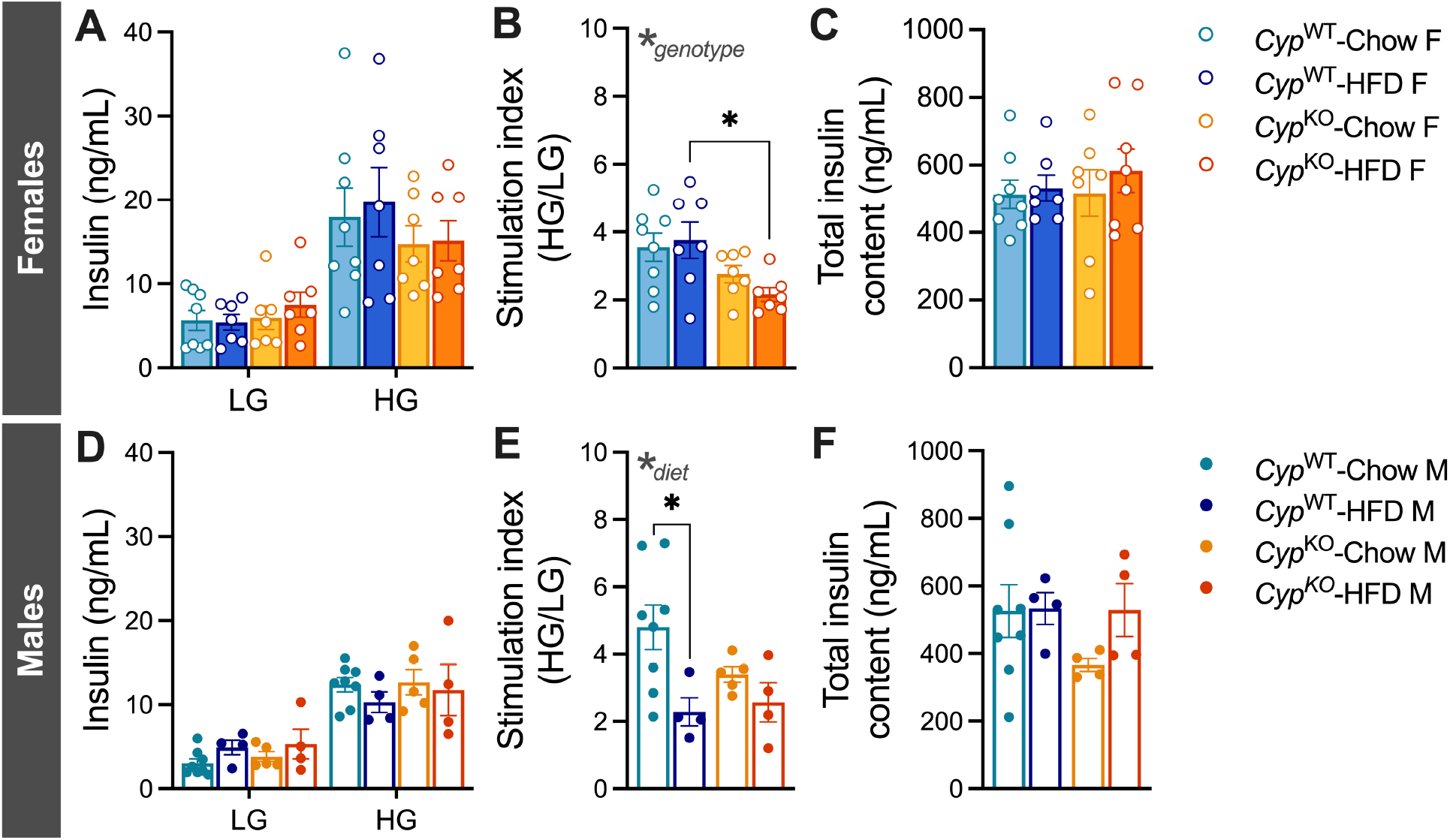
*Cyp1a1/1a2* are involved in regulating GSIS in both sexes. Islets were isolated at week 14 to assess glucose-stimulated insulin secretion (GSIS) *ex vivo* (see Figure 1A for the study timeline). LG = low glucose (2.8 mM glucose), HG = high glucose (16.7 mM glucose). (A, D) Insulin secretion following a sequential 1-hour incubation in LG and HG buffer in (A) female and (D) male islets. (B, E) Stimulation index in (B) female and male mice; stimulation index was calculated by dividing the average insulin concentration under HG by the average insulin concentration under LG for each biological replicate. (C, F) Total insulin content in islets of (C) female and (F) male mice following an overnight incubation in acid ethanol. There were 7-8 female mice and 4-8 male mice per group. All data are presented as mean ± SEM. Individual data points on bar graphs represent biological replicates (different mice). The following statistical tests were used: 2-way ANOVA with Tukey’s post-hoc test; *p < 0.05, **p < 0.01.

### 3.7 *Cyp1a1* induction in *Cyp^WT^*-HFD male islets is associated with increased expression of other stress response genes

Next, we assessed the expression of key genes related to stress response or beta cell function in islets isolated from *Cyp*^WT^ and *Cyp*^KO^ mice at the study endpoint. *Cyp1a1* and *Cyp1a2* expression were absent in islets from *Cyp*^KO^ compared to *Cyp*^WT^ females, as expected; HFD feeding did not change *Cyp1a1* or *Cyp1a2* expression in *Cyp*^WT^ female islets (**Figure 7Ai-ii**). *Nrf2*, a critical regulator of the oxidative stress response [37] **(Figure 7Aiii)***, Ucp2*, an uncoupling protein involved in regulating mitochondrial ROS production [38] **(Figure 7Av)**, and *Chop*, an indicator of endoplasmic reticulum stress [39] **(Figure 7Avi)** were unaffected by either diet or genotype in female islets, but diet had an overall effect to modestly increase *Gpx1*, a glutathione peroxidase involved in reducing ROS [40] **(Figure 7Aiv)**.

**Figure 7.**
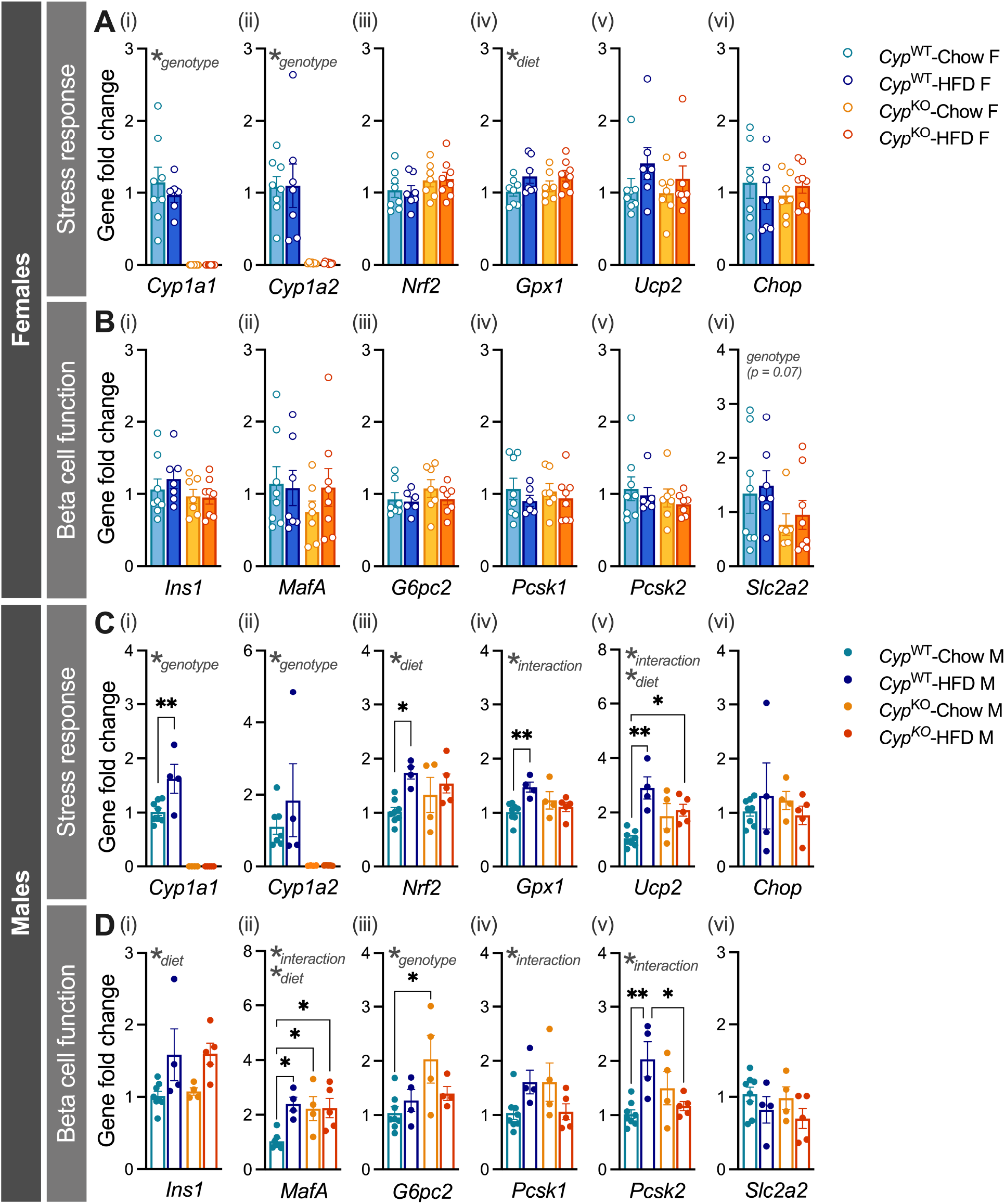
*Cyp1a1* induction in *Cyp^WT^*-HFD male islets is associated with increased expression of other stress response genes. Gene expression was measured by qPCR in islets isolated at week 14 (see Figure 1A for the study timeline). Expression of (A, C) genes involved in beta cell maturity, glucose uptake and metabolism, and insulin processing; (B, D) genes involved in the cellular stress response in (A, B) female (n = 6-8 mice per group) and (C, D) male islets (n = 4-8 mice per group). All data are presented as mean ± SEM. Individual data points on bar graphs represent biological replicates (different mice). 2-way ANOVA with Tukey’s post-hoc test; *p < 0.05, **p < 0.01.

Despite genotype-based differences in glucose homeostasis and stimulation index in females, levels of *Ins1* (insulin gene), *MafA* (transcription factor critical for maintaining the identity of mature beta cells [41]), *G6pc2* (gene involved in converting glucose-6-phosphate to glucose [42]), and *Pcsk1 and Pcsk2* (prohormone convertase genes required for converting proinsulin to insulin [43]) were unchanged by genotype or diet in female islets **(Figure 7Bi-v).** Only *Slc2a2*, a glucose transporter gene, showed a modest trending decrease due to genotype in female islets **(Figure 7Bvi)**.

Interestingly, HFD feeding increased *Cyp1a1* by *∼*1.6-fold in *Cyp*^WT^ male islets compared to chow-fed *Cyp*^WT^ male islets **(Figure Ci)**; *Cyp1a2* expression was not affected by diet **(Figure Cii)**. Islets from *Cyp*^WT^-HFD males also displayed an upregulation in *Nrf2* (∼1.7-fold), *Gpx1* (∼1.4-fold), and *Ucp2* (∼2.9-fold) compared to islets from *Cyp*^WT^-Chow males; these effects were absent in *Cyp^KO^*islets **(Figure 7Ciii-v)**. Islet expression of *Chop* was not affected by HFD feeding in either genotype **(Figure 7Cvi)**. Collectively, these results suggest that the induction of *Cyp1a1/1a2* is driving some of the stress responses in male islets.

Genes related to islet function were also differentially expressed in male islets. Levels of *MafA* **(Figure 7Dii)**, *Pcsk1* **(Figure 7Div)**, and *Pcsk2* **(Figure 7Dv)** were influenced by an interaction between diet and genotype. Diet had an overall effect on *Ins1* **(Figure 7Di)** and *MafA* **(Figure 7Dii)**, while genotype had an overall effect to increase *G6pc2* **(Figure 7Diii)**. Specifically, *MafA* was ∼2.4-fold higher in *Cyp*^WT^-HFD vs *Cyp*^WT^-Chow islets, whereas HFD did not increase *MafA* expression in *Cyp*^KO^ islets **(Figure 7Dii)**. *Pcsk2* was ∼2-fold higher in *Cyp*^WT^-HFD islets compared to *Cyp*^WT^-Chow islets, whereas this phenotype was abolished in *Cyp*^KO^ male islets **(Figure 7Dv)**. *Slc2a2* was unchanged by diet or genotype in male islets **(Figure 7Dvi)**.

Altogether, our results highlight sex-specific differences in the response of islets to HFD-induced stress at the transcript level. In female mice, neither diet nor genotype had a strong effect on the expression of stress response genes or islet function genes, but there were notable genotype-driven effects on islet gene expression in males.

## 4. Discussion

In this study, we investigated the role of CYP1A1/1A2 enzymes in maintaining glucose homeostasis under baseline conditions and in response to HFD feeding. Chow-fed *Cyp*^KO^ females had lower body weight and plasma insulin levels than their *Cyp*^WT^ counterparts. Despite having reduced plasma insulin levels, *Cyp*^KO^-Chow females had comparable glucose tolerance and insulin sensitivity to *Cyp*^WT^-Chow females. On HFD, *Cyp*^KO^ females developed milder glucose intolerance than *Cyp*^WT^ females. The metabolic phenotype of male mice differed substantially from females. *Cyp*^WT^ males became insulin resistant and hyperinsulinemic within 2 weeks of HFD feeding, whereas *Cyp*^KO^ males only developed hyperinsulinemia after 7 weeks of HFD feeding. Interestingly, HFD feeding downregulated *Cyp1a2* in liver from male and female *Cyp*^WT^ mice but upregulated *Cyp1a1* only in islets from male *Cyp*^WT^ mice. Elevated islet *Cyp1a1* levels were associated with increased expression of other stress markers in *Cyp*^WT^-HFD male islets, whereas HFD feeding did not alter the expression of these stress genes in islets from *Cyp*^KO^ males. While we observed genotype-driven differences in islet function and gene expression, liver and adipose phenotypes were primarily driven by diet, not genotype. Overall, we demonstrate that deleting *Cyp1a1/1a2* genes protects against HFD-induced glucose dysregulation in a sex-specific manner. Our results also underscore a potentially novel role for islet AhR signalling in regulating glucose homeostasis.

Several studies have shown that disrupting the AhR pathway improves metabolic outcomes in mice following HFD feeding [25–31], and studies that compare male and female mice reveal sex-specific effects [26, 27]. Our study further supports these findings by demonstrating that deleting AhR downstream targets, *Cyp1a1/1a2*, blunts the metabolic defects caused by HFD feeding in a sex-specific manner. We found that *Cyp*^KO^ females were generally leaner than *Cyp*^WT^ mice and, when fed HFD, *Cyp*^KO^ females developed milder glucose intolerance than *Cyp*^WT^ females. Similar results were found in global *Ahr* knockout (*Ahr*^KO^) female mice, where *Ahr*^KO^ females were protected from HFD-induced weight gain, exhibited higher energy expenditure, and had better glucose tolerance compared to *Ahr* wildtype (*Ahr*^WT^) females [27]. While plasma insulin levels were not assessed during the GTTs in the *Ahr*^KO^ study [27], our results show that chow-fed *Cyp*^KO^ females had lower plasma insulin levels than chow-fed *Cyp*^WT^ females. Furthermore, isolated *Cyp*^KO^ female islets also had reduced GSIS *ex vivo* compared to *Cyp*^WT^ female islets, pointing to a role for *Cyp1a1/1a2* in regulating insulin secretion and/or insulin clearance. Collectively, disrupting the AhR pathway, either by inhibiting AhR activation or preventing downstream *Cyp1a1/1a2* induction, is protective against HFD-induced weight gain and glucose dysregulation in female mice.

In contrast to female mice, there were fewer *in vivo* metabolic differences between *Cyp*^WT^ and *Cyp*^KO^ male mice. Males of both genotypes maintained similar body weight and glucose tolerance on either chow or HFD. However, *Cyp*^WT^ males exhibited HFD-induced hyperinsulinemia earlier than *Cyp*^KO^ males and became insulin resistant after only 2 weeks of HFD feeding. A similar pattern was observed in a global *Ahr*^KO^ model, where HFD-fed *Ahr*^KO^ males developed less severe insulin resistance than HFD-fed *Ahr*^WT^ males [27, 28]. Interestingly, *Ahr*^KO^ males were also partially protected from HFD-induced weight gain and glucose intolerance [27, 28]. Plasma insulin levels were not assessed in HFD-fed *Ahr*^KO^ males so the effect of *Ahr* deletion on HFD-induced hyperinsulinemia remains unclear [27, 28]. Altogether, the findings from the *Ahr*^KO^ studies and our *Cyp*^KO^ study suggest that in male mice, metabolic dysfunction from HFD feeding is partly mediated by AhR downstream targets, *Cyp1a1* and *Cyp1a2*.

To explore tissue-specific roles of *Cyp1a1/1a2* in driving the metabolic response to HFD feeding, we examined liver, fat, and islets, as these tissues are critical for maintaining glucose homeostasis. Given the central role of the liver in xenobiotic clearance, we hypothesized that deleting xenobiotic metabolism enzymes, *Cyp1a1/1a2*, will have the greatest impact on the liver. We also expected the effect to be pronounced in fat, since previous reports have shown that AhR activity is involved in HFD-induced obesity [25–31]. The role of the AhR pathway is least understood in islets, but we hypothesized islets would be highly susceptible to damage associated with CYP1A1/1A2 activity, given their relatively weak antioxidant defense system [44, 45]. Interestingly, HFD feeding induced *Cyp1a1* in islets of *Cyp*^WT^ male mice, but downregulated *Cyp1a2* in liver of both male and female *Cyp*^WT^ mice; neither *Cyp1a1* nor *Cyp1a2* was detectable in fat. Since CYP1A1 is the primary CYP1A isoform in non-liver tissues and CYP1A2 is the predominant isoform in the liver [46], tissue-specific variations in *Cyp1a1/1a2* regulation were expected. Our results align with previous reports showing reduced *Cyp1a2* gene expression and protein activity in rodent liver after HFD feeding [47, 48], but the induction of *Cyp1a1* in islets following HFD feeding has only been reported previously by our group [32]. Importantly, the induction of *Cyp1a1* in islets appeared to drive other stress responses in islets (e.g., *Gpx1* and *Ucp2* upregulation), as these changes were absent in *Cyp*^KO^ islets. In contrast, differences in liver pathology (i.e., steatosis, ballooning, inflammation) and adiposity were generally driven by HFD feeding but not genotype in both sexes. In all, these findings suggest that *Cyp1a1* induction in islets is an important physiological response to HFD feeding.

Previous reports have shown that exposure to TCDD, a strong chemical inducer of the AhR pathway and *Cyp1a1/1a2*, leads to reduced plasma insulin levels [49–51] and impaired *ex vivo* GSIS [49, 52, 53]. Given that *Cyp*^KO^ mice are unable to activate CYP1A1/1A2 enzymes, we expected *Cyp*^KO^ islets to have a more robust GSIS than *Cyp*^WT^ islets. Interestingly, chow-fed *Cyp*^KO^ females had lower plasma insulin levels than their *Cyp*^WT^ counterparts, and islets isolated from *Cyp*^KO^ females had lower GSIS than *Cyp*^WT^ female islets. Despite having less glucose-responsive islets, *Cyp*^KO^ females had similar glucose tolerance to *Cyp*^WT^ females on chow diet and better glucose tolerance than *Cyp*^WT^ females when fed HFD, but normal insulin sensitivity. The decreased insulin requirement in *Cyp*^KO^ females was associated with a trending decrease in the glucose transporter gene (*Slc2a2*) in *Cyp*^KO^ islets compared to *Cyp^WT^* islets, but not in liver (*Slc2a2*) or adipose (*Slc2a4*) tissues. Future studies should examine the role of *Cyp1a1/1a2* in insulin secretion and glucose uptake by beta cells.

In our study, deleting *Cyp1a1/1a2* genes provided greater protection for female than male mice from HFD-induced metabolic dysfunction, yet the underlying reason for this sex difference remains unclear. Rodent studies have shown that *Cyp1a1* expression (lung, kidneys) and CYP1A2 enzyme activity (liver) are higher in female tissues than in males [54, 55]. Therefore, the benefits of global deletion of *Cyp1a1/1a2* could be magnified in females because their baseline levels of CYP1A1/1A2 are higher than in males [54]. The interaction between hormones and CYP1A1/1A2 enzymes might also be involved, given that sex hormones (e.g., estrogen, progesterone) are endogenous substrates of CYP1A1 and CYP1A2 [46]. Estrogen levels may also contribute to female resilience against HFD-induced damage, as estrogens have a protective effect in islets and can even enhance beta cell function in rodents [56–58]. Furthermore, female islets exhibit greater resistance to endoplasmic reticulum stress than male islets [59], enabling them to respond more effectively to increased metabolic demands, such as higher insulin requirements, during HFD feeding. In all, our findings further emphasize the need to evaluate the influence of biological sex in metabolic studies.

The protective effect of deleting *Ahr* on HFD-induced metabolic dysfunction has been reported [24–28, 31], but the role of AhR downstream targets, *Cyp1a1/1a2*, remained unclear. Here, we provide evidence that inducible *Cyp1a1/1a2* genes may potentiate the harmful effects of HFD feeding on islets and systemic glucose homeostasis. Given that we used a global *Cyp*^KO^ model, we are unable to differentiate the role of CYP1A1/1A2 enzymes at the level of a beta cell, islet, or organism. Future studies should explore beta cell-specific or islet-specific knockout models to better elucidate the role of AhR and CYP1A1/1A2 in the endocrine pancreas.

## Supporting information

Ching et al., 2024 - supplemental file

## Abbreviations

AhR: Aryl hydrocarbon receptor
*Cyp1a1*: Cytochrome P450 1A1
*Cyp1a2*: Cytochrome P450 1A2
*Cyp*^KO^: Global *Cyp1a1/1a2* double knockout mice on C57Bl/6J background
*Cyp*^WT^: *Cyp1a1/1a2* wildtype C57Bl/6J mice
GSIS: Glucose-stimulated insulin secretion
GTT: Glucose tolerance test
HFD: High-fat diet
ITT: Insulin tolerance test
ROS: Reactive oxygen species

## 5. Acknowledgements

This research was supported by an NSERC Discovery Grant to J.E.B. (RGPIN-2017-06265) and a CIHR Operating Grant to A.M.C. (PJT-419982). J.E.B. was also supported by an Early Researcher Award from the Ontario Government. M.E.A.C. was supported by NSERC CGS-M and NSERC CGS-D awards. M.P.H. was supported by a CIHR CGS-D award. L.B. was supported by an CIRTN-R2FIC-CREATE and CIHR CGS-D award. E.v.Z. was supported by an CIRTN-R2FIC-CREATE award.

## 6. Author Contributions

M.E.A.C., M.P.H., and J.E.B. conceived the experimental design and analysed the data. M.E.A.C. and J.E.B. wrote the manuscript. M.E.A.C., M.P.H., L.B., and J.E.B. were involved with data collection and interpretation. R.M. and E.v.Z. were involved with data collection. A.M.C. assisted with tissue processing and liver pathology assessments. All authors contributed to manuscript revisions and approved the final version of the article.

