## Supplementary material for "CYP1A1/1A2 enzymes mediate glucose homeostasis and insulin secretion in mice in a sex-specific manner": Ching et al., 2024 - supplemental file

### Supplement Table 1. Primer sequences used for qPCR.

| Gene | Forward sequence | Reverse sequence | Amplicon size (bp) |
|---|---|---|---|
| ***Chop*** | TTGAAGATGAGCGGGTGGCA | CCAGGTTCTGCTTTCAGGTGT | 115 |
| ***Cyp1a1*** | ATCACAGACAGCCTCATTGAGC | AGATAGCAGTTGTGACTGTGTC | 139 |
| ***Cyp1a2*** | CAAGAGGTTTAAGACCTTCAATGATAAC | AAAGATGTCATTGACAATGTTGACAAT | 193 |
| ***G6pc2*** | AGCTGCCCTAAGCTACACCA | AACACTCCACAGAAAGGACCA | 85 |
| ***Gpx1*** | CTCACCCGCTCTTTACCTTC | CACACCGGAGACCAAATGATG | 103 |
| ***Ins1*** | TCAGAGACCATCAGCAAGCA | CTCCCAGAGGGCAAGCAG | 89 |
| ***MafA*** | AGTCGTGCCGCTTCAAG | CGCCAACTTCTCGTATTTCTCC | 149 |
| ***Nrf2*** | CTCTGCTGCAAGTAGCCTCG | TGTCTTGCCTCCAAAGGATGT | 138 |
| ***Pcsk1*** | GGTGGAAAGGTCGAGTCTAGC | TGCACACCAAACGCAAAAGA | 153 |
| ***Pcsk2*** | TTTGGAGTCCGAAAGCTCCC | GGTGTAGGCTGCGTCTTCTT | 91 |
| ***PPIA*** | GCCAGGACCTGTATGCTTTA | AGCTCTGAGCACTGGAGAGA | 178 |
| ***rRNA*** | AAACGGCTACCACATCCAAG | GCTGGAATTACCGCGGCT | 190 |
| ***Slc2a2*** | GCAACTGGGTCTGCAATTTT | CCAGCGAAGAGGAAGAACAC | 88 |
| ***Slc2a4*** | GCCCGGACCCTATACCCTAT | GTCACTCGCTGCCGAGG | 132 |
| ***Tnfα*** | AGTCCGGGCAGGTCTACTTT | ATGAACACCCATTCCCTTCA | 55 |
| ***Ucp2*** | CACCGGCAGCTTTGAAGAAC | GGGACCTTCAATCGGCAAG | 129 |

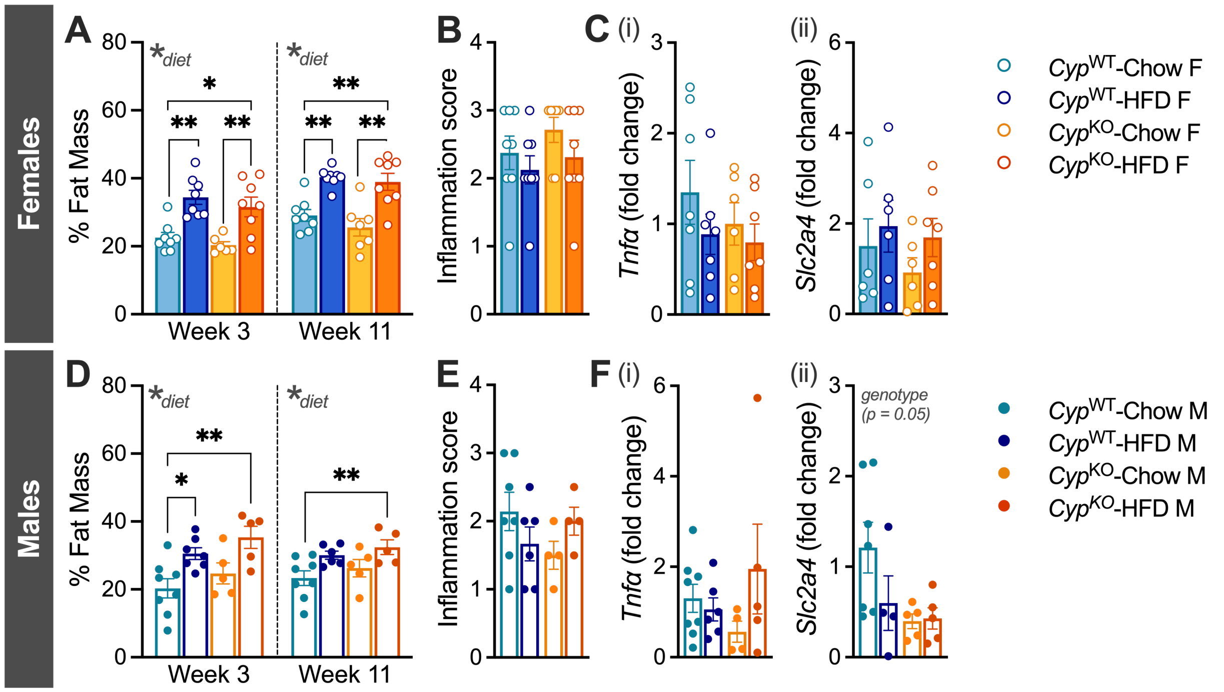


**Supplement Figure 1. Knocking out *Cyp1a1/1a2* did not result in significant changes in adiposity.** EchoMRI results (% fat mass) at weeks 3 and 11 (see Figure 1A for the study timeline) in (A) female (n = 7-8 per group) and (D) male mice (n = 5-8 per group). (B, E) Peripancreatic fat tissues collected at week 14 were stained with H&E and scored for inflammation (7-8 female mice per group, 4-8 male mice per group); scores ranged from 0 (no or minimal inflammation) to 3 (severe inflammation). (C, F) Transcript levels of the inflammatory marker, *Tnfα*, and the glucose transporter, *Slc2a2*, in peripancreatic fat at week 14 in (C) female (5-8 per group) and (F) male mice (n = 4-8 per group). The following statistical tests were used: 2-way ANOVA with Tukey's post-hoc test, *p < 0.05, **p < 0.01.
